# Constraints on eQTL fine mapping in the presence of multi-site local regulation of gene expression

**DOI:** 10.1101/084293

**Authors:** Biao Zeng, Luke R. Lloyd-Jones, Alexander Holloway, Urko M. Marigorta, Andres Metspalu, Grant W. Montgomery, Tonu Esko, Kenneth L. Brigham, Arshed A. Quyyumi, Youssef Idaghdour, Jian Yang, Peter M. Visscher, Joseph E. Powell, Greg Gibson

**Affiliations:** School of Biological Sciences and Center for Integrative Genomics, Georgia Institute of Technology, Atlanta, GA 30332, USA; Institute for Molecular Biosciences, University of Queensland, Brisbane, Australia; Estonian Genome Center, University of Tartu, Tartu, Estonia; Department of Medicine (Emeritus), Emory University, Atlanta, GA 30329, USA; Department of Medicine, Division of Cardiology, Emory University, Atlanta GA 30322, USA; Division of Biology, NYU Abu Dhabi, Saadiyat Island, United Arab Emirates

## Abstract

Expression QTL (eQTL) detection has emerged as an important tool for unravelling of the relationship between genetic risk factors and disease or clinical phenotypes. Most studies use single marker linear regression to discover primary signals, followed by sequential conditional modeling to detect secondary genetic variants affecting gene expression. However, this approach assumes that functional variants are sparsely distributed and that close linkage between them has little impact on estimation of their precise location and magnitude of effects. In this study, we address the prevalence of secondary signals and bias in estimation of their effects by performing multi-site linear regression on two large human cohort peripheral blood gene expression datasets (each greater than 2,500 samples) with accompanying whole genome genotypes, namely the CAGE compendium of Illumina microarray studies, and the Framingham Heart Study Affymetrix data. Stepwise conditional modeling demonstrates that multiple eQTL signals are present for ~40% of over 3500 eGenes in both datasets, and the number of loci with additional signals reduces by approximately two-thirds with each conditioning step. However, the concordance of specific signals between the two studies is only ~30%, indicating that expression profiling platform is a large source of variance in effect estimation. Furthermore, a series of simulation studies imply that in the presence of multi-site regulation, up to 10% of the secondary signals could be artefacts of incomplete tagging, and at least 5% but up to one quarter of credible intervals may not even include the causal site, which is thus mis-localized. Joint multi-site effect estimation recalibrates effect size estimates by just a small amount on average. Presumably similar conclusions apply to most types of quantitative trait. Given the strong empirical evidence that gene expression is commonly regulated by more than one variant, we conclude that the fine-mapping of causal variants needs to be adjusted for multi-site influences, as conditional estimates can be highly biased by interference among linked sites.

## Introduction

Since it is now recognized that the majority of SNP-trait associations identified by GWAS can be attributed to effects on gene expression, precise estimation of the location and effect sizes of regulatory polymorphisms has become important for understanding the relationship between genetic and phenotypic variation (Maurano et al. 2012; Farh et al. 2015). Expression quantitative trait locus (eQTL) analysis and related functional genomic strategies are thus now a standard component of genetic fine mapping (Nicolae et al. 2010). The minimal expectation is that they can identify the gene within a locus that accounts for a GWAS signal, although even this is a far from trivial undertaking (Chung et al. 2014; Pickrell 2014). Many investigators make the stronger assumption that co-localization of eSNP and GWAS signals to a tight linkage disequilibrium interval implies the ability to define if not the causal variant, then at least a credible set of SNPs that include the causal site (Trynka et al. 2013; Gaulton et al. 2015; Kichaev and Pasaniuc 2015; Liu et al. 2016). The strong enrichment of chromatin marks such as DNAse Hypersensitive Sites (DHS) in the vicinity of eQTL validates this assumption (ENCODE Project Consortium 2012; Roadmap Epigenomics Consortium 2015), but conversely raises the question of why there are so many instances of discordant fine localization: does this reflect biochemistry (regulatory sites need not map to ENCODE elements), or simply limits to the statistical resolution of association signals (Udler et al. 2010)?

The general and parsimonious assumption is that functional variants are sparsely distributed, and hence that their precise localization or estimation of effect sizes is not affected by interference due to confounding of statistical signals. However, as genome-wide association studies have increased in size it has become clear that multi-site effects are not uncommon. For example, the latest meta-analysis of height suggests that over one third of the more than 400 identified loci have multiple independent signals (Wood et al. 2014), and the expression of a large proportion of genes in lymphocyte cell lines is regulated by two or more locally-acting variants (cis-eQTL) (Liang et al. 2013). Since linkage disequilibrium within a locus can be extensive, the potential for mis-estimation of eQTL effects due to interference between signals from tightly linked polymorphisms is high. Here we address this concern through a combination of simulation studies and empirical analysis of two large peripheral blood gene expression cohort studies with whole genome genotype data.

Heritability analyses in recent years have shown that on average up to half of the variance of phenotypic traits, or of transcript abundance, can be explained by genetic factors, mostly acting in an additive manner (Powell et al. 2013; Wright et al. 2014). An important difference with visible phenotypic traits is that one or a few SNPs are often found to explain a large proportion of the genetic variance. These typically lie within 1Mb of the transcript and are regarded as cis-acting regulatory polymorphisms. As shown in a companion study (Lloyd-Jones LR et al. 2016) of the CAGE dataset, 35% of all expressed genes in peripheral blood have narrow sense heritability greater than 0.1, with a median of 0.3. The sentinel cis-eQTL typically explains 85% of the locally acting variance, which is two thirds of that attributed to all detected eQTL, but the majority of the genetic variance is generally actually due to trans-acting polymorphisms of small effect.

The largest blood eQTL study reported to date, assembled from meta-analysis of over 5,000 individual Illumina microarray samples (Westra et al. 2013), reports single site local associations that are genome-wide significant for 6,418 genes (44% of those tested) with a 5% false discovery rate. However, the blood eQTL browser only provides single-site (unconditional) estimates for all local SNPs at each locus. A more powerful cross-population Bayesian method (Gusev et al. 2015) applied to just 420 lymphocyte cell lines in the Geuvadis dataset (Lappalainen et al. 2013) found a very similar number of eGenes, 14% of which had strong evidence for secondary association signals in a multi-site analysis. The primary aim of our study was to combine both of these strategies by performing comprehensive conditional eQTL mega-analysis of two large predominantly European-ancestry cohorts: the Consortium for the Architecture of Gene Expression (CAGE), consisting of ~2,800 Illumina microarray samples from 5 studies conducted in Brisbane, Atlanta, Estonia, and Morocco (Idaghdour et al. 2010; Powell et al. 2012; Kim et al. 2014; Schramm et al. 2014; Wingo and Gibson 2015), and the Framingham Heart Study consisting of ~5,000 Affymetrix samples from the offspring and third generations of the FHS (Huan et al. 2015). Imputed whole genome genotypes were available for both cohorts.

While increased sample size is certain to reveal more specific instances of multi-site regulation, it also provides the opportunity to more accurately define effect sizes in the presence of multiple sites that have varying degrees of linkage disequilibrium. Figure 1 illustrates the reasoning for a hypothetical case. Five true eSNP effects are indicated by red lines with increasing effects above the horizontal and decreasing below it, while single site unconditional estimates at common variants across the locus are indicated as black lines. Where two sites in high LD have effects in the opposite direction (green circle), they will either cancel each other out, or substantially bias the effect size estimates. Where two sites act in the same direction (blue circle), their effects will tend to be added together and hence the strongest association will over-estimate the effect while the secondary site will be under-estimated or not detected. Weaker associations (brown circle) may also go un-detected if they are influenced by even low levels of linkage disequilibrium. In theory, if the location of the functional sites is known *a priori*, these difficulties can be resolved by multi-site linear regression simultaneously fitting all of the identified SNPs. In practice, the identities of the functional sites are unknown, and exhaustive multi-site modeling is impractical, so sequential conditional analyses are used to find secondary, tertiary, and so forth associations that are independent of the primary signal. In the presence of strong LD, this approach is expected to miss independent associations which will remain confounded with the primary signal.

**Figure 1.**
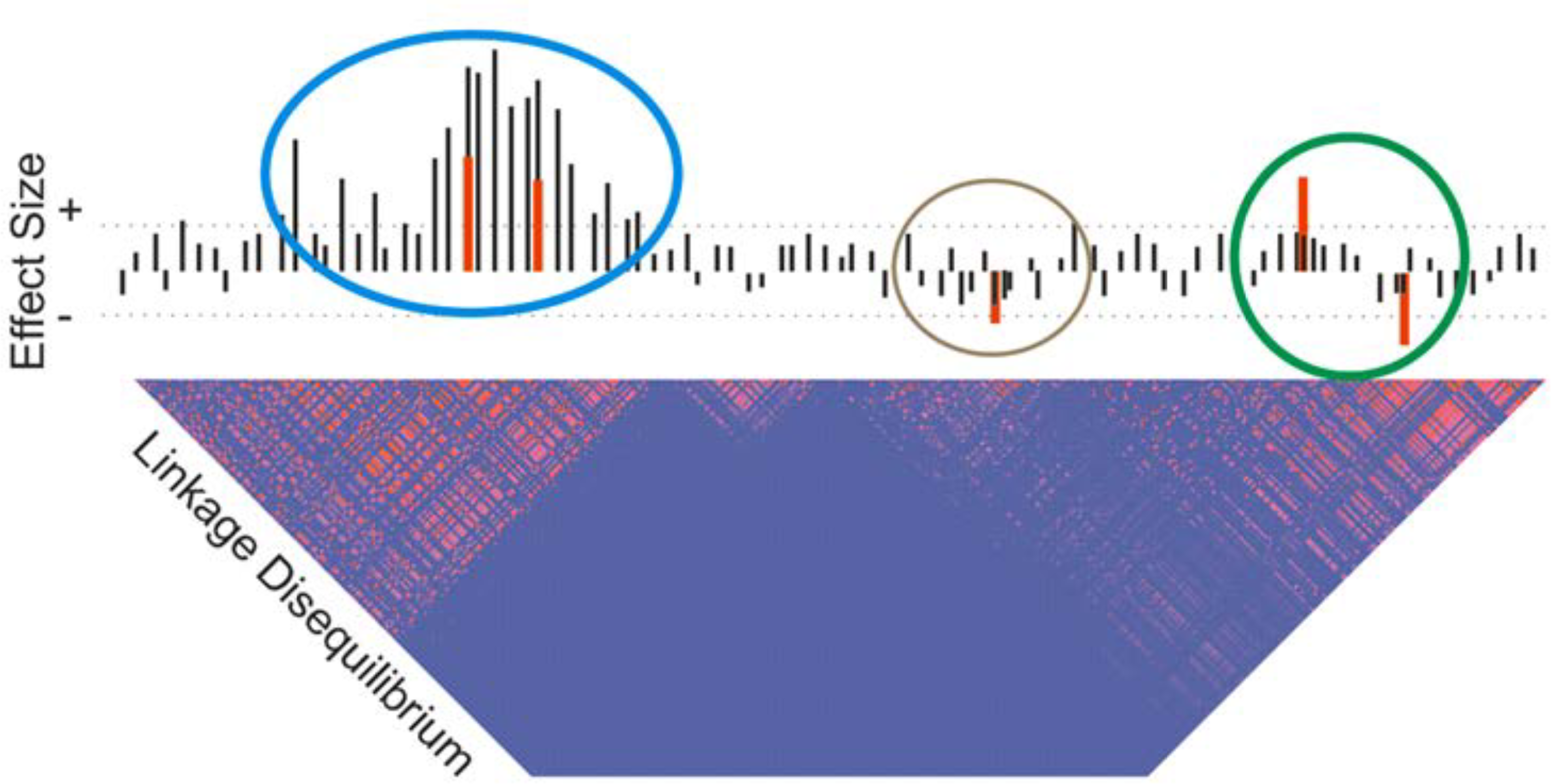
Schematic of multi-site regulation of gene expression. Black bars indicate univariate estimates of allelic effects of minor alleles increasing (above the horizontal) or decreasing (below the horizontal) gene expression without conditioning on other sites. Red bars show the actual effects at 5 SNPs in this locus, which has a linkage disequilibrium profile with two large and one small block of elevated LD (pink squares). Dotted horizontal lines indicate a statistical significance threshold, which is only exceeded in the univariate modeling by the two left-hand sites (blue circle). Since these two sites act in the same direction, they reinforce one another, leading to over-estimation of their effect sizes, whereas the two at the right (green circle) interfere with one another antagonistically, leading to underestimation of their effects. The effect at the fifth site (brown circle) may only be identified following conditional analysis.

This study explores the sources of error in estimating eQTL effects. The design and workflow is summarized in Figure S1. Empirical analysis of the CAGE and FHS datasets finds that statistical evidence for 2, 3 or even more sites regulating expression of a gene is common, although the concordance in localization of signals to tight LD intervals lower than expected. We then use simulations to explore the effect of untagged variants and multi-site regulation on the localization of, and effect size estimation of statistical peaks, concluding that there is likely to be a meaningful fraction (between 5% and 25%) of all discovered eQTL credible intervals that do not actually include the causal variant.

## Results

### 1. Multiple eQTL regulation is ubiquitous in human blood

Our primary objective was to estimate the proportion of loci that have multi-site regulation in the two large cohort studies, CAGE and FHS. Since both datasets include siblings, we used GEMMA (Zhou and Stephens 2012) to perform sequential conditional eQTL analysis deploying a genetic relationship matrix based on all measured and imputed genotypes to model family structure. Before evaluating the real data, we assessed the performance of this strategy by simulation, using the package SeqSIMLA2 (Chung et al. 2015) to simulate family-based data. Figure S2 shows that in the absence of LD, the conditional approach underestimates the allelic effects, likely because a portion of the genotype effects are absorbed into the genetic relatedness matrix, whereas the estimated effect sizes with the multilocus model centered at the expected values. Similarly, in the presence of intermediate LD, the multilocus method was less biased, indicating that sequential modeling followed by joint fitting of discovered allelic effects is a reasonable approach to multiple eQTL discovery.

Applying sequential conditional analysis to CAGE, we detected 5,974 eGenes (37.8% of 15,812 tested genes) with at least one significant eSNP at P < 10^−5^. Of these eGenes, 2,187 (36.6%) contain probes influenced by more than one eSNP and hence appear to be regulated by multiple enhancers. (Note that in the case of genes with multiple probes on the Illumina platform, we only required that at least one probe was associated with an eQTL, and for multi-SNP regulation included only the probe with the largest number of significant independent eSNPs). Similarly, in the FHS data, we detected 5,597 (35.3% of 15,853 tested genes), 2,098 (37.5%) of which were regulated by multiple eQTLs. In CAGE, the average variance explained by detected eSNPs was 6.1%, the same as in FHS, 6.1%, and in both cases these estimates account for more than half of the previous estimated *cis* heritability. For those genes with multiple eQTL regulation in CAGE, which have a mean explained variance of 7.2%, the newly detected secondary eSNPs typically explained 20% more variance than the peak SNP alone, namely ~1.2% of the phenotypic variance (6.0% vs 7.2%), in line with estimates from the companion study (Lloyd-Jones et al. 2016). For eGenes with multiple eQTLs in FHS, the mean explained variance is 6.3%, and the secondary signals increase the explained variance from 6.5% to 8.3%, an ~28% increase.

Figure 2 shows frequency histograms for the number of detected eQTL per gene in both studies: the number of loci with additional independent sites reduces by approximately two thirds with each additional SNP in both CAGE and FHS, up to half a dozen variants, and a few loci have 10 sites. This reduction likely reflects the true prevalence of multi-site effects as well as reduced power to detect SNPs that explain less of the variance than the primary signal. A detailed example of multi-site association is shown for the *HBZ* locus in CAGE (Figure 3), where from left to right, and top to bottom are the results of stepwise conditional analysis yielding 9 independent eQTL signals. The total explained variance is 39.8%, one third more than the 28.4% explained by the highest single-site signal. An example from the FHS is *ABHD2* where we detect 5 independent eQTLs explaining 9.3% of the variance, compared with 5.6% for the peak eSNP (although the Affymetrix probeset contains a common variant, rs2283435, that is in linkage equilibrium with each of the five regulatory signals). All of our multiple eQTL results can be downloaded as locuszoom figures from our lab server, and are provided as Table S2.

**Figure 2.**
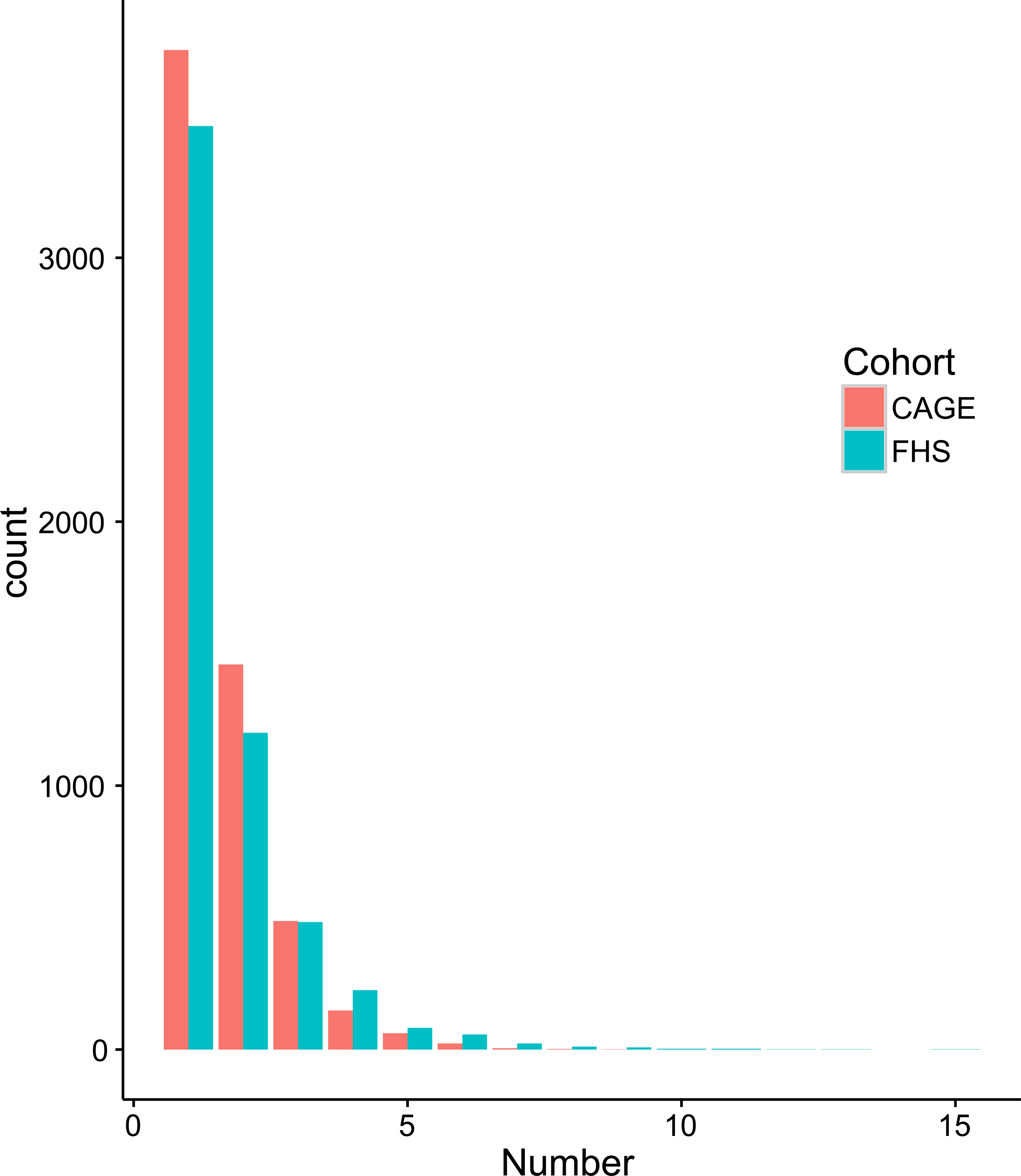
Number of detected eSNPs for each probe in CAGE and FHS cohorts. There are 16,467 Illumina probes expressed in whole blood across the CAGE studies (red), and 15,853 Affymetrix probesets for the FHS (green). Bars indicate the number of eGenes with the indicated number of independent local eSNP associations at p<10^−5^ following the sequential conditional analysis.

**Figure 3.**
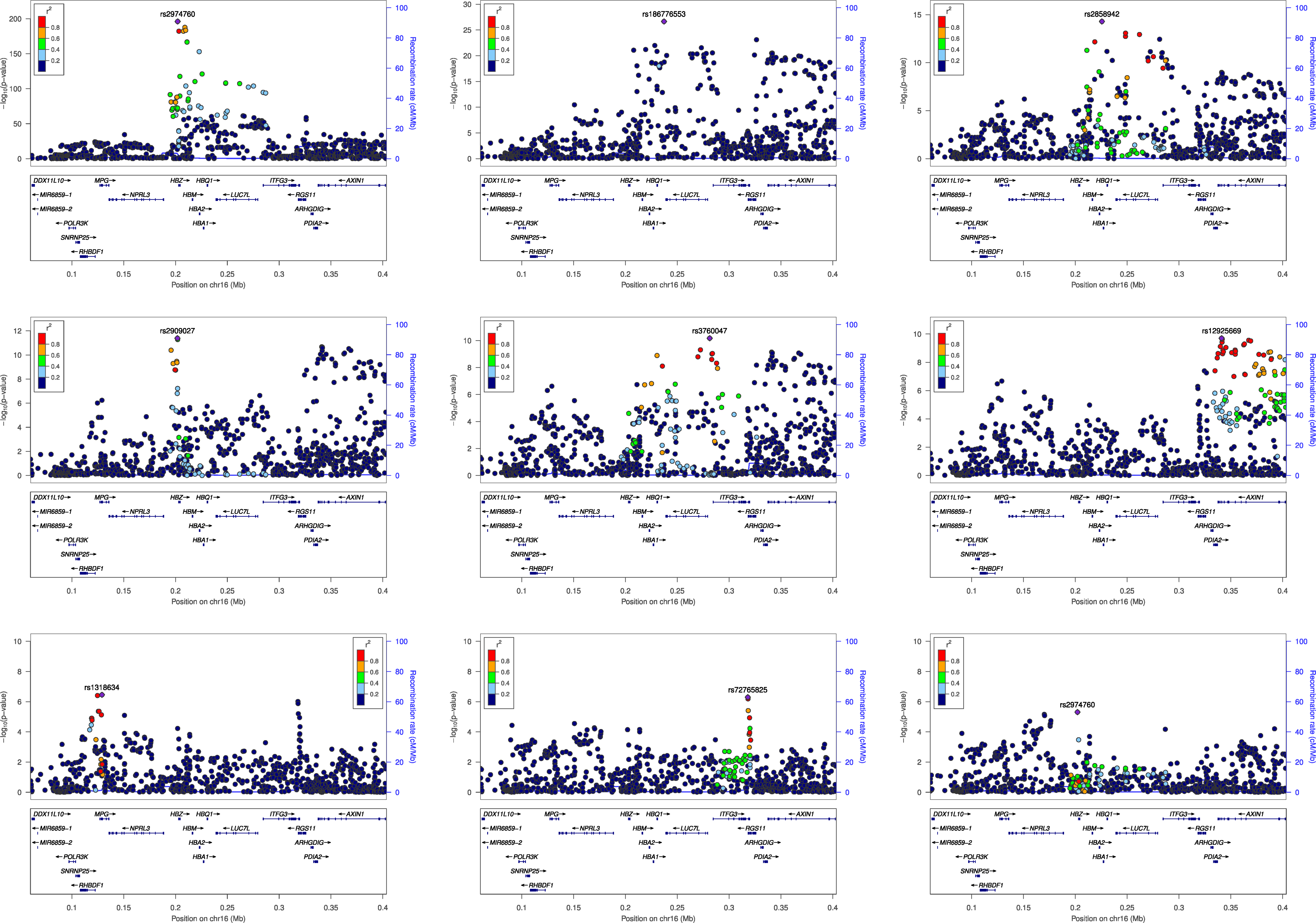
Example of multi-site association with gene expression. Each locus zoom plot shows the univariate association statistic (negative log p-value) for all variants in vicinity of the *HBZ* locus in the CAGE dataset, following each successive round of conditional analysis on the residuals fitting the previously identified peak SNP. Sites in linkage disequilibrium with the new peak SNP are colored according to the *r*^2^ value. The *HBZ* probe, ILMN_1713458, does not contain any known common variants.

We also computed the difference between the estimates fitting all discovered variants jointly and the conditional single-site estimates following eSNP sequential conditional discovery. The average change in estimated beta was 0.04 sdu, plus or minus 0.06 due to a long tail of large deviations. We have no way of directly evaluating whether the multi-locus estimates are more accurate than the conditional ones, but Figure 4A shows a heat map of differences between the two models. By comparison with the simulated data, three features are noteworthy. First, in the domain where multi-site modeling results in the strongest improvements in simulated data discussed below (Figures 4D and 4E), for SNPs of moderate to large effect in high LD, the joint modeling tends to revise effect size estimate downwards by up to a standard deviation unit (blue hue: negative values indicate larger effect estimates in single site models). Second, this is not apparent for SNPs in low LD. Third, for small effects in regions of high LD, the joint modeling tends to increase effect size estimates, also by up to a standard deviation (yellow hue).

**Figure 4.**
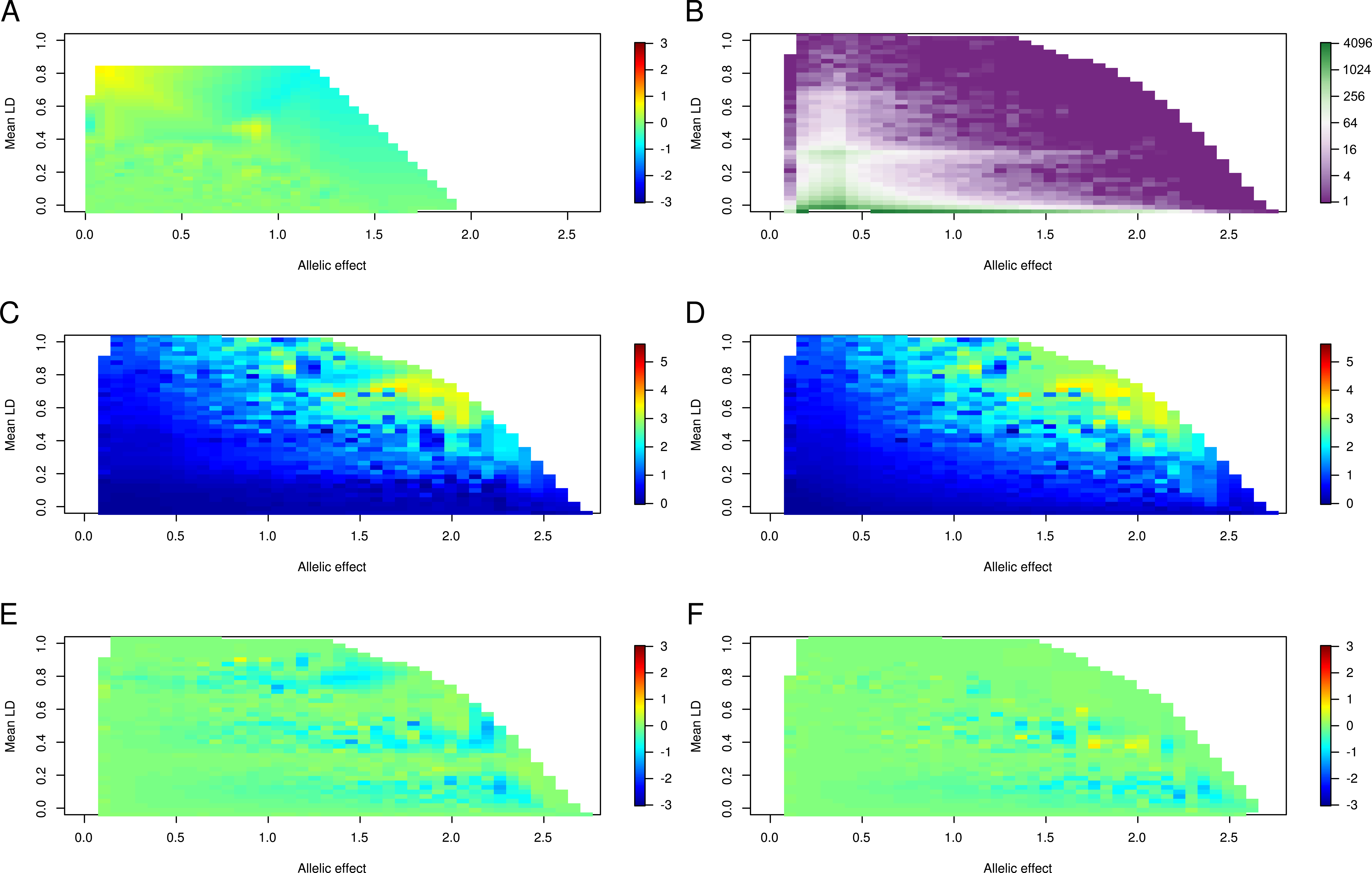
Biases in effect size estimation from conditional and joint analysis. Panel A compares estimates from joint and conditional modeling in the CAGE dataset, as a heatmap of the average difference where yellow indicates that joint modeling produces a larger estimated effect size, and blue a lower estimate. The remaining panels refer to 500,000 simulated data points where effect sizes were sampled from a uniform distribution to explain from 2% to 10% of the expression at a locus for each of 4 SNPs picked at random from 400kb intervals of the CAGE genotype data. Panel B shows the density distribution on the log2 scale of the number of simulations with alleles for each pixel with the indicated β (in standard deviation units, sdu) on the x-axis, and average LD with the other 3 sites at the locus on the y-axis. Panels C and D show the average absolute value of the deviation between the observed and known effect size for sites under the multi-site model where all discovered sites are fit jointly (C) or from single site estimates after each step of sequential conditional analysis (D), for the 4:0 scenario where all minor alleles have effects in the same direction. Panel E is the average difference between these two models (joint minus sequential), showing three bands of negative values indicating greater bias in the conditional estimated. Panel F is the difference between the models under the 3:1 scenario (see Figure S7 for the equivalent of C,D and for the 2:2 scenario).

The comparison of results from the CAGE and FHS analyses suggests a disappointingly low level of replication. The overall overlap between CAGE and FHS for eSNPs defining credible intervals with LD *r*^2^>0.8 is ^~^29.1%. Primary peaks in CAGE were detected for 53.0% of the eGenes represented in the FHS, and reciprocally 56.5% of the FHS eGenes had primary signals in CAGE, very similar to the proportions reported for eSNPs at P < 10^−8^ across four peripheral blood studies (Zeller et al. 2010) that had a variety of technical differences. Furthermore, only 41.5% of the primary signals in FHS are in LD (*r*^2^>0.8) with the primary eSNP in CAGE, suggesting different largest-effect regulatory variants are tagged in the two datasets. This overall eSNP replication rate is slightly higher (47.2%) when mapping to the transcript level for 1,314 probesets which map directly to the same exon and have an eQTL signal on both platforms. The replication rate of secondary, tertiary, and quaternary signals in FHS is 19.0%, 11.3%, and 11.0%, implying a successive decay. The reasons for the discrepancies may have to do with collapsing of probe-level data down to gene-level signals losing information on splice isoforms, the different normalization strategies (which alone can double discovery rates (Qin et al. 2012)), and cross-study heterogeneity. Comparison of the percent variance explained by discovered SNPs on the two platforms in Figure S3 shows that many genes are disproportionately tagged by eQTL in the two studies, implying platform effects. Among 139 genes with more than 20% of the variance explained by cis-eSNPs, the replication rates are 64.7% for the primary signal, 34.8% for the secondary, 14.4% for the tertiary, and 8.5% for the quaternary. The subsequent panels confirm that all of these replication rates are proportional to the percent variance explained overall, arguing that statistical power is also a major source of low replication.

In HT12 v3 Illumina probes, 10% of the uniquely mapping probes contain at least one SNP with MAF>1% in 1000 genome European population. The prevalence of eQTL was twice as great for these probes (59% versus 30% of 23,681 “clean” probes), so we just removed the Illumina probes containing SNPs in order to control the false discovery rate. However, since most of the Affymetrix probesets contain at least one SNP, this was not practical for the FHS dataset and instead we employed a conditional analysis strategy incorporating SNPs in probes as covariates. For ~15,000 detected eSNPs, one third of the association signals were abrogated by conditioning on the SNPs in probes, and the number of eGenes correspondingly reduced by 25%. Table 1 contrasts the eQTL results from both platforms before and after controlling for the SNP-in-probe effects. The first three data columns show the cumulative number of eGenes with at least 1, 2, 3, 4, or more detected eSNPs before SNP-in-probe removal, and the next three show the cumulative numbers after. The proportion of overlapping signals is not greatly affected. The last column shows that the number of eGenes where at least one detected signal is likely capturing the same variant is around 44% (689/1565), and that the number where all of the multiply detected signals are within *r*^2^ > 0.8 is very small. There are 214 genes with at least two signals in high LD with one another.

**Table 1.**
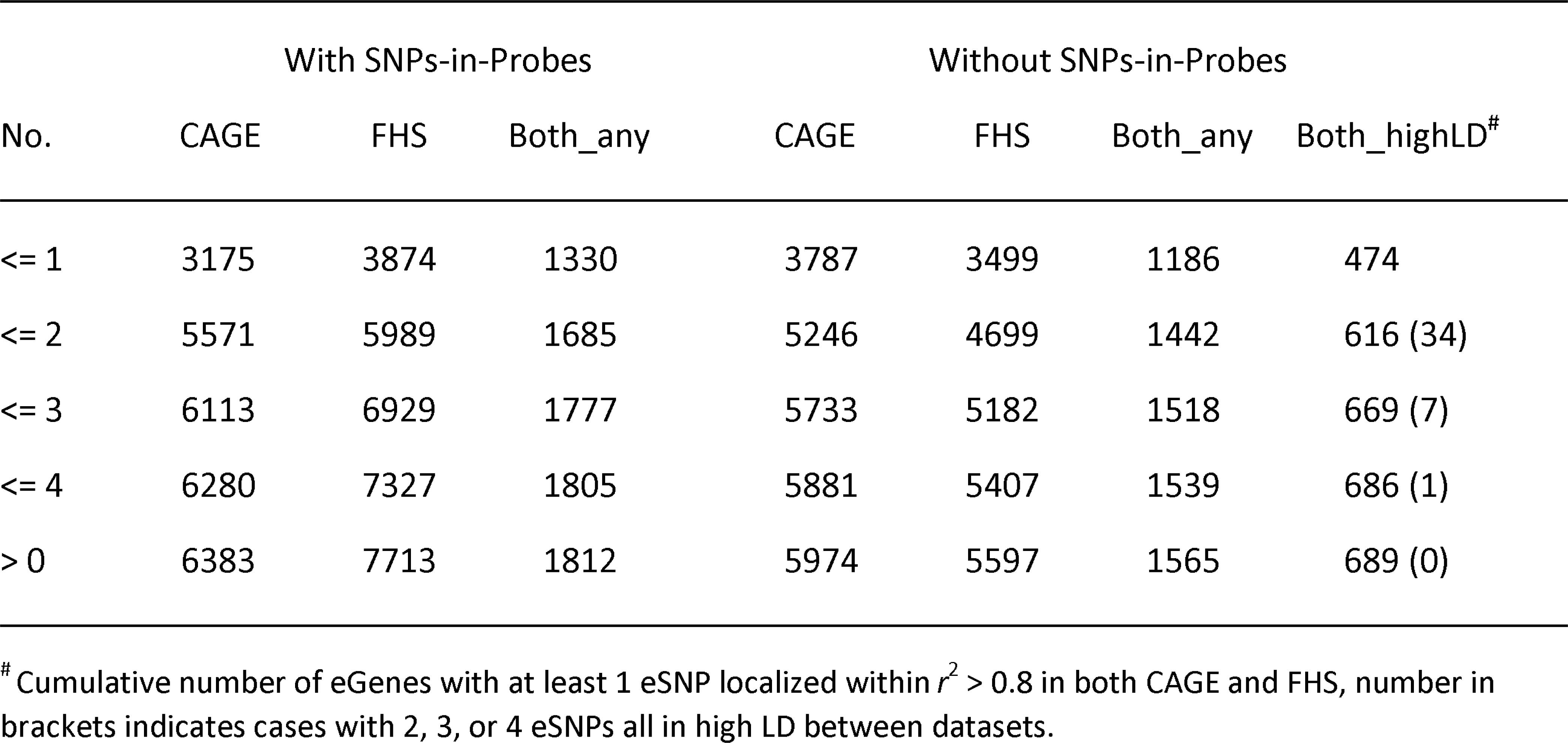
Cross-platform comparison of eSNP detection after adjustment for probe SNPs

| No. | With SNPs-in-Probes |  |  | Without SNPs-in-Probes |  |  |  |
| --- | --- | --- | --- | --- | --- | --- | --- |
|  | CAGE | FHS | Both_any | CAGE | FHS | Both_any | Both_highLD<sup>#</sup> |
| <= 1 | 3175 | 3874 | 1330 | 3787 | 3499 | 1186 | 474 |
| <= 2 | 5571 | 5989 | 1685 | 5246 | 4699 | 1442 | 616 (34) |
| <= 3 | 6113 | 6929 | 1777 | 5733 | 5182 | 1518 | 669 (7) |
| <= 4 | 6280 | 7327 | 1805 | 5881 | 5407 | 1539 | 686 (1) |
| > 0 | 6383 | 7713 | 1812 | 5974 | 5597 | 1565 | 689 (0) |

<sup>#</sup> Cumulative number of eGenes with at least 1 eSNP localized within $r^2 > 0.8$ in both CAGE and FHS, number in brackets indicates cases with 2, 3, or 4 eSNPs all in high LD between datasets.

### 2. Up to 10 percent of secondary associations may be false positives

Although these simulations indicate a high prevalence of multisite regulation of transcript abundance, some fraction of the events might be attributed to splitting of the effect due to a single imperfectly tagged site into two or more spurious signals. An example is shown in Figure 5A and 5B, contrasting the local Manhattan profiles for a single causal variant that splits into two associations when the site is excluded from analysis. To explore the frequency with which this occurs, we conducted simulations assuming a single causal variant based on the genotypes measured in 1,839 CAGE samples (excluding the BSGS samples, to avoid complications due to the presence of twins). Randomly assigning causal effects to sites genotyped in CAGE resulted in the appearance of a secondary signal at P <10^−5^, conditioned on the causal site, at 0.3% of the loci. This is approximately as expected given 8.3 million imputed SNPs at 22,000 loci, and is also the same as the false discovery rate of primary signals in the absence of any simulated causal locus. That is to say, our random expectation is for 0.3% of transcripts to have a false eQTL discovery at P <10^−5^ in the CAGE dataset.

**Figure 5.**
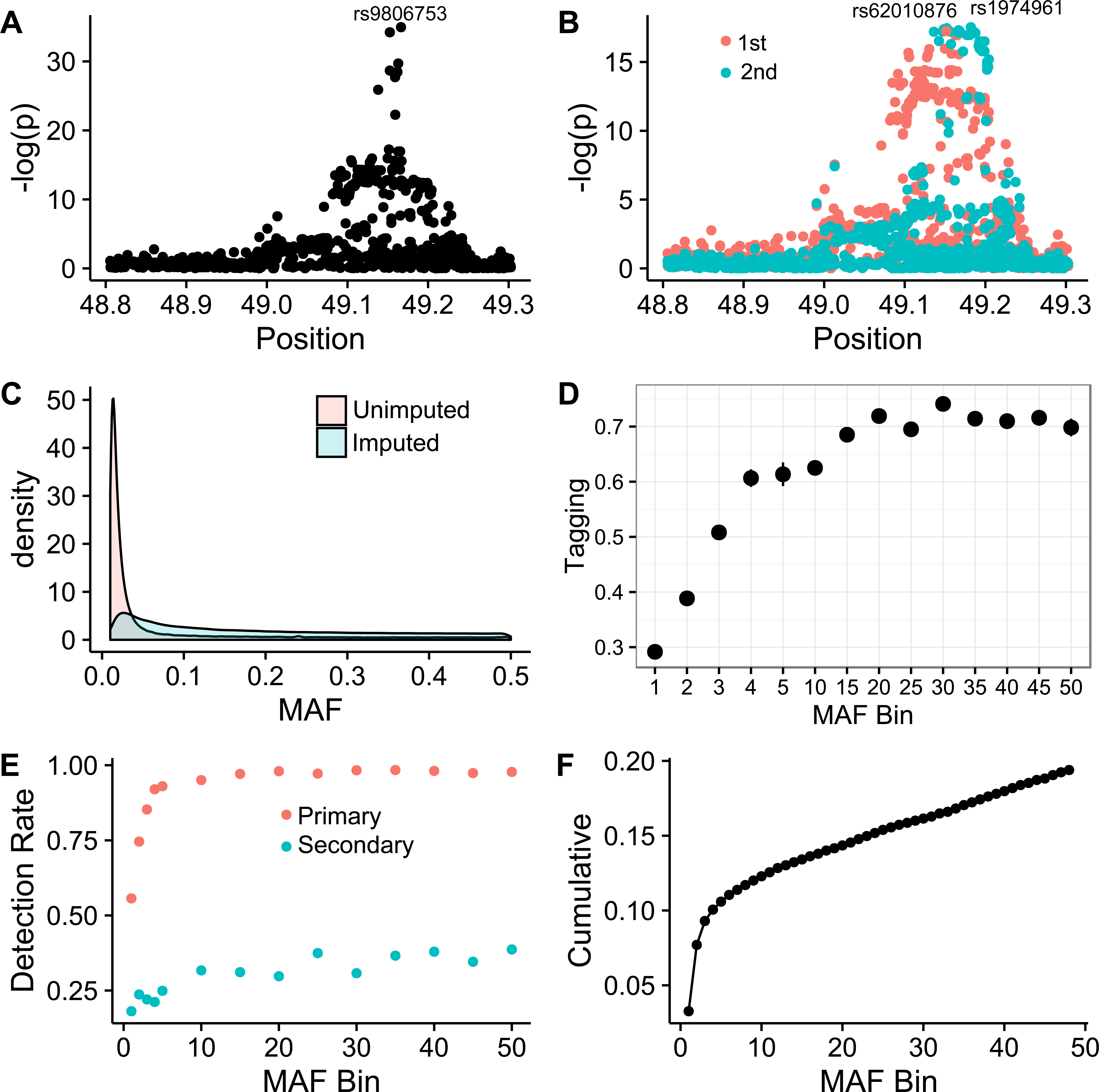
Proportion of false multiple eQTL detection due to unimputed variants. (A, B) Example showing how poor tagging can split one causal into two separate signals at the *SHC4* locus. (A) SNP rs9806753 (European ancestry maf = 0.23) was simulated in CAGE to generated an eSNP effect, but removal of this variant and all SNPs within *r*^2^ > 0.5 from the analysis (B) results in the effect being captured by both a primary (rs62010876, maf = 0.10) and secondary (rs1974961, maf = 0.13) signals. (C) Empirical maf density distribution of 1.4M unimputed 1000G (red) and 8.3 M imputed 1000G variants in CAGE (blue), demonstrating shift to lower frequencies for variants not tagged. (D) Tagging efficiency as a function of maf based in mean *r*^2^ for the strongest correlated SNP for 10,000 randomly selected variants across the frequency spectrum in the CAGE genotype dataset. (E) Corresponding signal detection rate at p<10^−5^ for randomly assigned effect sizes explaining between 2% and 8% of a simulated gene expression trait, for primary (red) and secondary (blue) signals when the simulated variant is excluded from the analysis. (F) Cumulative proportion of sites expected to generate a false multiple eQTL detection, calculated as the sum of the false secondary signal detection rate weighted by the maf frequency in the indicated bins. Multiplication of this proportion by the number of unimputed SNPs in genic regions (14% of 1.4M) and the actual proportion of SNPs that have effects (unlikely to be >1%) yields up to 400 possible false positive secondary associations.

However, this ignores the possibility that the causal variants are not present in the imputed genotypes. Of the 9.7 million SNPs, indels and CNV with maf > 0.01 in the European populations in the 1000G database, 1.9 million are not imputed in CAGE. Figure 5C shows that the minor allele frequency distribution for these variants is strongly shifted toward rarer alleles relative to the imputed SNPs, and is centered at a maf ~ 0.02. Consistent with Yang et al. (2015), the average tagging efficiency (*r*^2^ value) of these SNPs is a function of minor allele frequency, being greater than 0.7 for maf > 0.05, but dropping to less than 0.5 for maf = 0.01, as seen in Figure 5D.

Since we cannot simulate effects at non-imputed SNPs, we approximated such alleles by randomly simulating a causal variant from the CAGE SNPs with the same frequency distribution, but excluding it from the analysis along with all variants that would tag it at the typical level observed for the non-imputed SNPs of the same maf. We then asked how often the effect is captured by multi-site signals, as a function of residual tagging efficiency. We allowed for increased effect sizes with lower maf by simulating effects in the constant range of 2% to 10% of the variance explained. The proportion of such pseudo-unimputed SNPs which generate primary signals is reduced with lower maf, due in part to the smaller proportion of variance explained by the less common variants that partially tag them. For common variants, there is almost always a second site in high enough LD to capture most of the causal signal in the absence of genotypes at the causal variant, but rare variants are insufficiently tagged to generate a signal at all, 90% of the time.

In the presence of tagging SNPs with *r*^2^ > 0.5 to the “unobserved” causal variant, false secondary associations are observed up to 50% of the time. At the other end of the spectrum, rare variants (maf ~0.01) that produce a primary signal at a tagging SNP with 0.1 < *r*^2^ < 0.3 also produce a secondary signal but less frequently. The blue curve in Figure 5E indicates the inferred fraction of unimputed variants that could induce secondary signals as a function of maf, and Figure 5F shows that the cumulative proportion of such spurious eQTL weighted by observed maf proportions is approximately 20%. Approximately 14% of the 1.9M unimputed variants are located within 200 kb of a gene, and assuming that 0.5% of these actually have an eQTL effect, this suggests the potential for ~250 such effects.

These computations argue that up to 10% of the observed >2,000 multi-site associations have the potential to be false signals driven by inefficient tagging of unimputed variants in CAGE. The proportion could be greater if the fraction of functional SNPs is higher, as suggested for example by Tewhey et al. (2016) who used a very sensitive MPR assay to implicate 3% of regulatory sites in 3,642 eQTL regions (842/32,373 tests) as capable of modulating transcript abundance. However, the proportion of sites with detectable signals capable of explaining more than 2% of the variance is certainly lower, and even 1 in 200 (0.5%) is likely an over-estimate given that there are of the order of 1,300 documented variants in the vicinity of each gene: 6 eSNPs per gene seems to be an upper limit. Synthetic associations due to even more rare variants may be expected to generate split associations as well (Dickson et al. 2010; Zhu et al. 2012). Yang et al (2015) found that approximately 20% of the variance for height can be explained by SNPs with maf < 0.1, in part due to larger effect sizes of prevalent very rare SNPs, many of which are likely secondary associations. We also found that there is an excess of rare variants (maf <0.01) influencing extremes of gene expression, also with a slightly larger distribution of effect sizes than common variants (Zhao et al. 2016). Too many unknown parameters need to be evaluated to give a good estimate of the number of false positive secondary associations due to synthetic effects of very rare alleles, but it may be another few percent.

### 3. Multi-site effect estimates have variable accuracy, but may be improved by joint fitting

Our conclusion from the preceding section is that no more than 10% of all secondary associations with gene expression may be false positives due to split tagging of un-imputed variants. The next question is how accurate are the effect size estimates and localization of the SNPs in the remaining 90% of cases of true multi-site regulation likely to be? In this section, we describe simulations designed to evaluate (i) what proportion of eSNPs are expected to be detected by the sequential conditional approach; (ii) how large is the bias in effect sizes of the discovered SNPs; and (iii) how much the existence of multi-site regulation influences the fine-mapping of causal sites within credible intervals. These results can be used as a baseline for comparison with the empirical results from the CAGE and FHS studies presented above.

#### 3.1. Effect of joint fitting on effect size estimation for pairs of known causal variants

As a prelude to this analysis, it is worth noting that in the case where the identities of two causal variants are known *a priori*, joint fitting of the two SNPs in a single regression on transcript abundance always results in more accurate effect size estimates given a large sample size. The bias in estimation due to linkage disequilibrium between pairs of SNPs is a function of the two effect sizes (β_1_ and β_2_), the correlation between the SNPs (*r*), and the ratio of the square root of the product of their allele frequencies: 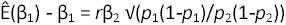 (Yang et al. 2012). This is maximized for pairs of SNPs at the same frequency, increases with high LD, and can be either positive or negative depending on whether the signs of the minor allele effects are coupled or not. Supplementary Figure S4 provides a visual summary of the biases, compared with the effect of jointly fitting the two SNPs with a sample size of 2,000, which uniformly improves the effect size estimates.

Three results deserve highlighting. First, for each combination of allele frequencies, increasing the allelic effect size results in more severe biases, in the most severe cases over-or under-estimating the effects by as much as 50% of the variance explained. Second, the first picked SNP (with the larger effect size) has the greater deviation between the estimated and true effect size. This makes intuitive sense as the larger effect will generally be the first detected one and absorbs much of the effect of the other SNP, which will typically be under-estimated in the conditional analysis, but to a lesser extent. If the deviation is computed simply on the uni-site unconditional values, the opposite result is obtained: the deviation is greatest for the smaller effect site. Third, the mis-estimation is greatest for lower allele frequencies, which is particularly noteworthy since most eQTL have maf in the range of 0.1 to 0.3.

We also considered the power to detect joint effects in the presence of LD. As the p value cutoff for detection becomes more stringent, the bias in estimation becomes more severe, since the first picked SNP absorbs the effect of both alleles into the same estimate, leaving the statistical power of the other allele, conditioned upon the first one, close to zero. With a sample size of 2,000 and intermediate LD, when both alleles are modeled jointly, power to detect both effects remains high across the plausible parameter space once the effect size exceeds 5% variance explained, and the estimation for the beta value is still accurate. Down-sampling suggests that in order to estimate effects within 0.1 standard deviation units, for pairs of variants with LD *r*^2^ ~ 0.9, each explaining 10% of the variance (namely having effect sizes of at least a half a standard deviation unit), a sample size of at least 900 is required.

Consequently, most small sample eQTL studies will fail to resolve linked sites into two effects. These results indicate how the typical assumption that an eQTL effect is due to a single variant in a set of credible SNPs in high LD is potentially highly biased. Similar conclusions apply to the situation where two SNPs operate in opposite directions, with the additional dilemma that they will not be detected at all and consequently strongly under-estimate the regulatory variance at a locus.

#### 3.2. Under-estimation of allelic effects by sequential conditional analysis

However, in most analyses, the identities of the causal variants are not known, and must be estimated from the data. Although Bayesian methods are being developed that have some promise for simultaneous detection of multi-site effects (Zhou et al. 2013), the current standard is to perform sequential conditional analysis, in which the residuals from discovery of each SNP are taken forward as the dependent variable in a new scan for an independent SNP (Yang et al. 2012). To explore the performance of this strategy in the context of four causal regulatory variants per transcript, 500,000 simulations were performed by randomly picking 4 SNPs within 200 kb up-or down-stream of a gene, from the genotypes of 1,839 unrelated individuals in the CAGE cohort. We assigned each SNP an allelic effect size so as to explain between 2 and 10% of the variance of a trait otherwise uniformly distributed with a mean of zero and standard deviation of 1. Power to detect individual effects of this magnitude is close to 100% at the significance level P < 10^−5^. The sampling was performed across all genes so as to sample from the typical LD structure in the European ancestry human genome, and four variants were chosen since the empirical evidence indicates that up to 10% of all loci may have such complexity. We also report the simulation results for 2 and 3 causal variants in Supplementary Figures S5 through S7. Furthermore, effects were randomly assigned under three scenarios, with either 4 positive (4:0), 3 positive and 1 negative (3:1), or 2 positive and 2 negative (2:2) effects on the trait. For the eQTL detection, once the sequential conditional detection was completed, we determined which of the four causal variants was in high LD (*r*^2^ > 0.8) with one of the discovered sites. If a peak was in high LD with more than one causal variant, it was assumed that it tagged the highest effect site.

Table 2 summarizes the “tagging efficiency”, namely the percent of simulations in which the indicated number of significant independent sites was detected, as well as the proportion of the discovered variants that are in high LD (*r*^2^ > 0.8) with one of the simulated causal variants. The proportion of credible intervals that contain the simulated site ranges between 85% and 90%. Notably, 10% of the simulations detected more than 5 independent peaks, at least one of which is spurious. Similar results are reported for the reciprocal measure of what fraction of simulated variants is captured by discovered variants in Table S3. Across the 500,000 simulations, the power to detect multiple eSNPs is a function of the number of sites operating in the same direction. It is highest for the case where the minor alleles for all four variants have effects in the same direction, and least where two are in one direction and two in the opposite direction. In the 4:0 scenario, 3 or more of the 4 eSNPs are detected three quarters of the time, whereas this proportion drops to two thirds the simulations for 2:2. No variants are detected in just over 1% of the simulations, and just one variant in approximately 6% of them, in most cases tagging a true causal variant. However, as the number of detected variants increases, so does the probability that one of them is not within *r*^2^ > 0.8 of one of the simulated causal variants.

**Table 2.** Tagging Efficiency of Detection of Causal Variants with *r*^2^ cutoff 0.8

| Number of<br>Detected Causal | Scenario (Positive: Negative effects)* |  |  |
| --- | --- | --- | --- |
|  | 4:0 | 3:1 | 2:2 |
| 1 | 0.6% (0.84) | 1.4% (1.00) | 1.2% (0.62) |
| 2 | 5.7% (0.69) | 10.4% (0.91) | 12.8% (0.86) |
| 3 | 28.2% (0.78) | 24.3% (0.85) | 22.5% (0.76) |
| 4 | 55.5% (0.89) | 59.0% (0.92) | 60.2% (0.90) |
| >4 | 10.0% (0.63) | 5.0% (0.66) | 3.3% (0.56) |

\* % indicates the percent of cases with the indicated number of independent discovered variants; numbers in brackets are the proportion of discovered variants that are in LD ( $r^2 > 0.8$ ) with one of the simulated causal variants.

Another way to consider the power of multi-site detection is to ask how much of the variance explained by the four SNPs is captured by the discovered variants. Box and whisker plots in Figure 6 show that under all three scenarios, on average 85-90% of the variance is captured, namely in these simulations approximately 15% to 20% of the transcript abundance. Although effect sizes of all SNPs were drawn from the same distribution, the first discovered SNP (rightmost box in each panel) typically explains between one third and one half of the expected variance, suggesting that it often tags some of the effect of another site, though this fraction is lower than that observed in actual data. As expected, summation of the independent contributions from the sequential conditional models, or fitting all of the discovered variants simultaneously, explains very similar proportions of the variance overall. Similar results are seen with 2 or 3 simulated causal variants (Figures S5 and S7).

**Figure 6.**
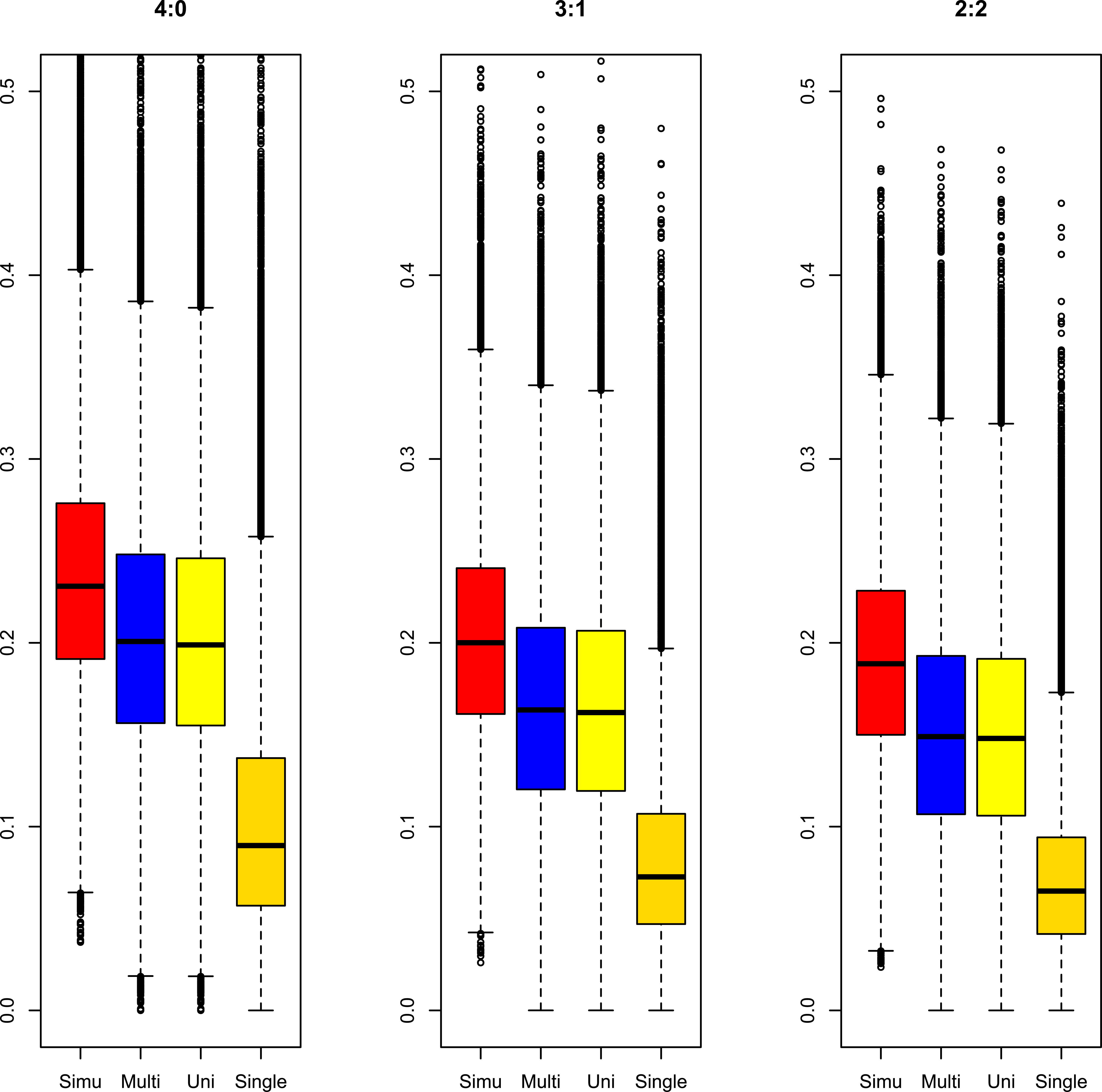
Proportion of variance explained by detected eSNPs in simulations. Box and whiskers show median, inter-quartile range, and 95% confidence limits for the proportion of variance explained under three scenarios for 500,000 simulations of 4 sites affecting gene expression. From left to right in each simulation, Simu is the variance explained by the known sites, Multi is the result fitting discovered eSNPs jointly, Uni is the result of summing the effects from sequential conditional modeling, and Single is the effect of the peak detected eSNP. Y-axis is the proportion of variance explained. Scenarios are 4:0, all four minor alleles with effects in same direction; 3:1, one minor allele effect in the opposite direction; 2:2, two minor alleles on one direction and the other two in the opposite direction.

#### 3.3. Mis-estimation of allelic effects sizes by sequential conditional analysis

Even though the sequential conditional and multi-site models capture essentially equivalent proportions of the variance tallied across sites, we expected there to be biases in estimation of individual site effects that would be reduced by the multi-site modeling. To quantify this difference, we computed the deviation between the observed and true simulated effect sizes (β in standard deviation units) for each discovered peak located within *r*^2^ > 0.8 of an independent causal variant, and evaluated the absolute value of these deviations as a function of the mean LD between the causal variant SNP and the other 3 causal variants in the model. Figure 4C and D show the average absolute value of the deviation for SNPs with the indicated true effect size and LD, for the sequential conditional and multisite models respectively, for the 4:0 scenario where all minor alleles operate in the same direction. Figure 4E shows the average difference in the absolute value of the estimates (multi-site deviation minus sequential conditional deviation), with negative values indicating that the sequential univariate model under-or over-estimates the effect size by a larger amount. The sampling density is illustrated in Figure 4B, while Figure 4F shows the average difference for the 3:1 model (the actual deviations for this model and the 2:2 model are provided in Figure S6). Figure S7 summarizes analogous simulations with just 2 or 3 causal variants.

Several results are noteworthy. First, for causal variants in low LD, as expected, neither model results in appreciable estimate bias, but once the average LD rises above 0.5, effects can be misestimated by more than the effect size. For example, for β = 0.5, the absolute value of the difference between the observed and true effect is typically between 1 and 2 s.d. units, which depending on the allele frequency may correspond to at least 2 percent of the total gene expression variance. Second, for large effect alleles, the mis-estimation is appreciable even at intermediate levels of LD, and it is not unusual for estimates to be off by as much as 4 s.d. units under either model. Third, overall, the multisite modeling corrects some of the sequential conditional analysis bias. Panel 5E suggests three bluish-tinged bands of results with LD centered around 0.1, 0.4, and 0.8 where the average difference between the models of the absolute differences in measured and true effect sizes is in favor of the multi-site model for allelic effects up to β = 0.5, becoming twice as great for very large effect alleles. The latter only account for a very small fraction of all simulated alleles. Fourth, similar trends are seen for the 3:1 scenario where one of the minor alleles operated in the opposite direction to the other three, as well as in the 2:2 scenario. Fifth, simulations with just 2 or 3 causal variants yield similar conclusions. The advantage of joint modeling is reduced in the presence of opposing allelic effects, but still prevalent in the region with high allelic effect size and low LD; and as noted above a large proportion of causal sites are not discovered in the 2:2 scenario, so are not included in the estimation.

In summary, stepwise conditional eQTL discovery is expected to discover between 70% and 80% of eQTL within realistic effect size ranges typical of those reported in the literature. Once discovered, multi-locus estimation of effect sizes provides slightly more accurate estimates than the estimates from sequential conditional models, but for large effect alleles in high LD the corrections can be substantial. The frequency distribution of the improvement in estimation provides a baseline for evaluation of the influence of multi-locus modeling as applied to the real datasets in Section 1 (Figure 4A).

#### 3.4. Bias in estimation of the causal variant location

Another, possibly more important, measure of estimation bias is the location of the peak SNP relative to the causal site. Several methods have been developed to evaluate the likelihood that a peak eSNP and peak GWAS signal is due to the same variant, including PICS (Farh et al. 2015) and CoLoc (Giambartolomei et al. 2014). Although not as statistically rigorous, the Regulatory Trait Concordance (RTC) score (Nica et al. 2010) is nevertheless intuitive and easily implemented on the scale of our simulations. It is essentially a ranking of the significance of the detected eQTL p-value relative to the casual site, RTC = (*N*_*SNPs*_ - *Rank*_causal SNP_)/*N*_*SNPs*_ where a value of 1 indicates identity, and 0 that the two sites are in the same locus but highly unlikely to be capturing the same signal. Figure 7 contrasts the RTC scores for the primary eQTL signals relative to the largest effect causal variant in each simulation, for the 4:0, 3:1, and 2:2 scenarios. For comparison, under a single variant model, RTC is always close to 1 as expected (note that since we do not simulate the GWAS signal as well, these values are inflated relative to data where the identity of the actual causal variant is unknown (Nica et al. 2010)). For 10% of simulations in the presence of multiple regulatory variants, the RTC score of the primary SNP drops below 0.9, again with greater tendency toward mis-estimation of the eQTL location in models with opposing effect directions of the minor alleles. This analysis confirms the results in Table 2 indicating that, up to 15% of all detected SNPs are not in high LD (*r*^2^ > 0.8) with a simulated variant in the imputed panel of SNPs.

**Figure 7.**
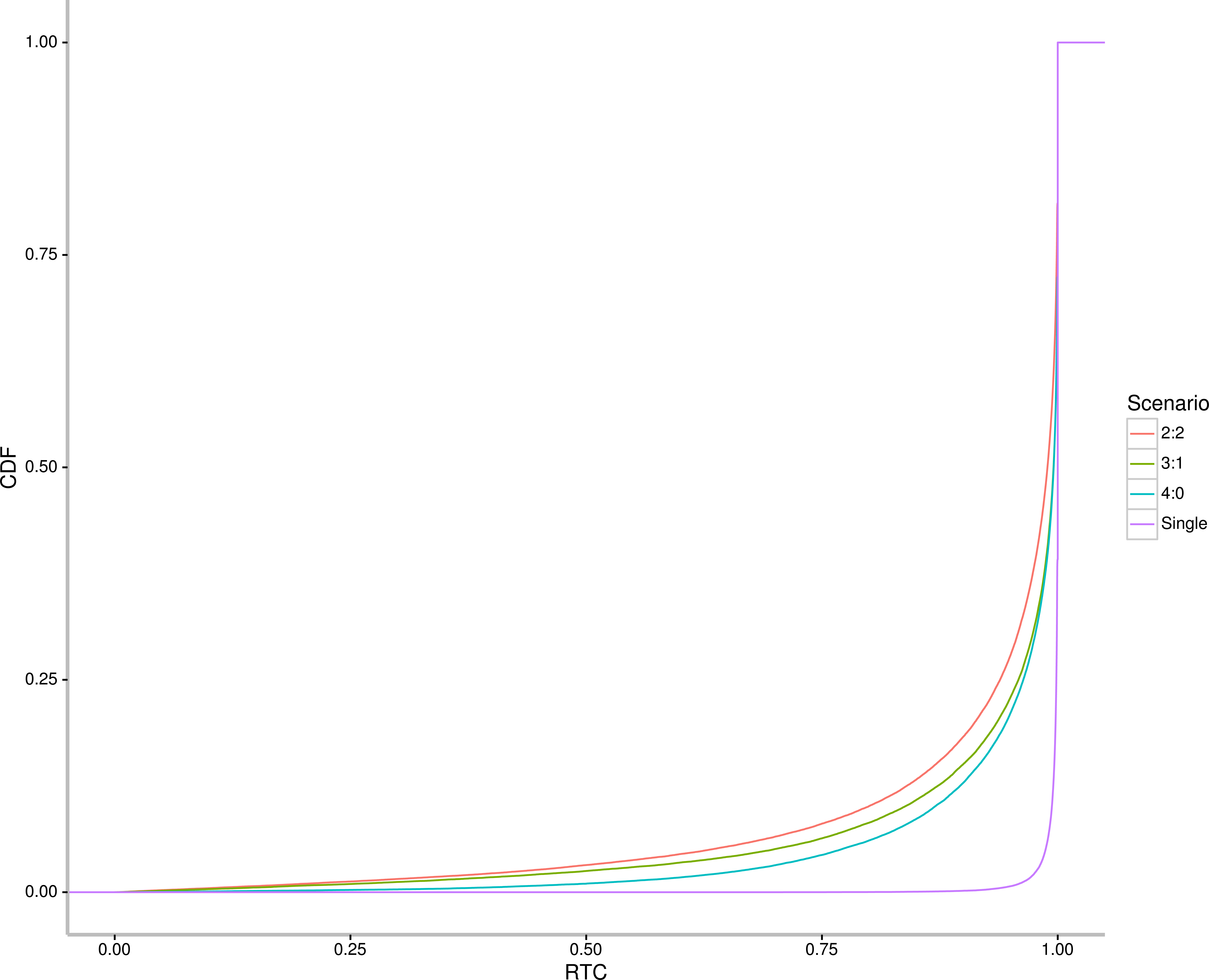
The single causal variant assumption biases fine mapping of causal variant locations. Each curve represents the cumulative probability distribution for RTC scores for the primary causal variants under a model with a single causal variant (purple), or with 4 causal variants under scenarios 4:0, 3:1, and 2:2 (blue, green, red curves respectively). RTC scores close to 1 imply equivalence of the significance values of the eQTL and causal variant.

Localization of tertiary and quaternary signals is affected more strongly, but intriguingly, considering just associations within *r*^2^ > 0.8 of a simulated SNP, the secondary signal is slightly more likely to be the first or second ranked SNP for one of the causal variants than is the primary signal. This is true under all three scenarios, which have very similar profiles to that shown for the 3:1 scenario in Figure S8 (since there is wide variance in the number of SNPs in each region, we simplified the analysis by reporting just the SNP ranks in this Figure, rather than RTC). It should be noted that there is not strong concordance between the relative proportion of variance explained by the causal variants and whether they are the primary through quaternary association, since LD has a strong influence on detection power. Although the vast majority of discovered sites are within 3 or 4 SNPs of at least one of the four causal variants *when they are in high LD with one of them*, it cannot be concluded that the order of discovery corresponds to the true order of effect sizes.

## Discussion

Studies of the genetic regulation of gene expression are making a meaningful contribution to the interpretation of GWAS results, as they provide functional insight into the nature of the causal genes. Efforts to fine map causal variants are, however, complicated by the limits of statistical resolution as it is not uncommon for tens, if not hundreds, of polymorphisms in a credible set to have similar statistical support (Gaulton et al. 2015; Kichaev and Pasaniuc 2015). Inclusion of experimental evidence from epigenetic marks or signatures of evolutionary conservation into scores such as CADD ((Kircher et al. and CATO (Maurano et al. 2015) promises to improve resolution, as do methods such as RTC (Nica et al. 2010) and PICS (Farh et al. 2015) which prioritize variants based on the structure of linkage disequilibrium at a locus. In general these approaches assume parsimony, namely that there is a single variant that is responsible for the major GWAS or eQTL signature. Although it has become increasingly clear that many loci harbor multiple independent regulatory variants, we argue here that if the parsimony assumption is relaxed and it is assumed that multiple sites in strong linkage disequilibrium commonly account for a signature that is compounded into a single significant association, then the estimates from sequential conditional analysis can be highly biased. To summarize, we find that over 5% of primary sites and more than a quarter of all causal sites are unlikely to be tagged at all; that in the presence of multi-site regulation at least 15% of all mapped sites are not in strong LD with any of the multiple imputed causal variants at a locus; and that another 10% of the associations are plausibly due to splitting of the signal due to an unimputed site. Taken together with the confirmation that over a third of all eGenes have two or more independent eSNPS, these results suggest that at least 5% and perhaps as many as a quarter of mapped credible intervals may not include the actual causal variant.

Theory and simulation both indicate that if two linked sites both influence a trait, including gene expression, then multi-site models will uniformly outperform sequential uni-site ones with regard to estimation of the true effect size. When the identities of the variants are known, sample sizes of several thousand individuals are sufficient to jointly estimate their effects with high accuracy even in the presence of high levels of linkage disequilibrium with r^2^ up to or even exceeding 0.9. The problem is that the identities of the variants are generally not known, and there are no established methods for comprehensive screening transcriptome-wide for localization of multi-locus local eQTL effects. Two exhaustive search algorithms, PAINTOR (Kichaev et al. 2014) and CAVIARBF (Chen et al. 2015) hold promise for detailed dissection of multi-site models at individual loci, and a Bayesian shotgun stochastic search algorithm, FINEMAP (Benner et al. 2016) has recently been proposed for rapid maximum likelihood estimation of multi-SNP contributions. This and a deterministic approximation of posteriors (DAP) approach to iterative refinement of multi-SNP models (Wen et al. 2016) should be evaluated for their ability to identify and estimate more of the multi-site effects than those obtained with the sequential conditional approach. We also caution that it is likely that single site effects may sometimes be artificially split into two or more linked contributions under each of these strategies. Consequently, for this study we adopted the existing standard approach of sequential conditional analysis. As recently reported by others (Gusev et al. 2015; Wen et al. 2016), a substantial number of loci have evidence for multiple such eSNPs at a locus, but our large sample sizes allow us to increase the estimated fraction to approximately 14% of all expressed transcripts in peripheral blood having at least two independent eSNPs, and ~10% having three or more, up to 9 sites for a handful of loci.

We then estimated the bias in the estimates from the conditional analysis, by fitting multi-locus linear models to all of the discovered eSNPs at each locus. This revealed only modest improvements in accuracy for most of the discovered sites, but the modesty is in part an artefact of the discovery bias introduced by the sequential conditional process. Our simulations assuming 2 to 4 effective sites per locus across a wide and representative range of linkage disequilibrium, show that in a sample size of 2,000, in general no more than 85% of the simulated causal sites are tagged by discovered associations that explain typically observed magnitudes of effect and would almost always be detected if a single site explained the variance. Similarly, Lloyd Jones et al. (2016) estimate that in the CAGE dataset, on average between 50% and 75% of the heritability due to locally acting regulatory polymorphism can be attributed to discovered variants. Multi-site modeling re-adjusts the remaining estimates typically by between 0.1 and 0.5 standard deviation units, which depending on the allele frequency accounts for between 2% and 5% of the variance explained, and only rarely more than 10%.

However, any variants with effect sizes greater than 1 sdu, and whose average *r*^2^ with the other 3 SNPs is greater than 0.9, will be mis-estimated in both the single site and joint models, typically by 1.5 sdu or more. The mis-estimation is on average the greatest where all of the effects are in the same direction, but is consistently observed also in the presence of associations with alternate signs. There is no way to know what proportion of the variants in the CAGE and FHS datasets are mis-estimated, but our analyses suggest that it is similar to the simulated data, with a non-negligible source of error stemming from the failure to separate single site effects into two or three contributing SNP effects.

Since the number of loci with 2, 3, 4 and so forth discovered variants approximately halves for each additional variant, it seems reasonable to infer that at least 10% of loci actually have four or more eSNPs (two or three of which are discovered), and that perhaps one quarter have three or more eSNPs (of which one or two are discovered). We conclude that the sequential conditional estimates of eQTL effects are actually highly biased for a considerable proportion of variants. Although this does not impact the total amount of variance explained by the discovered variants, it is likely to greatly impact fine mapping efforts, particularly where two or more effects are collapsed into one site in a credible interval.

One difference between the actual data and simulations is that the secondary SNPs explain substantially less of the variance at each locus. Figure 6 shows that the primary signal is only responsible for between one third and one half of the gene expression variance at each locus in the simulations, whereas they often account for more than half of the variance in actual whole blood. Similarly, LD score regression argues that primary SNPs typically capture more than half the variance for gene expression (Kichaev and Pasaniuc 2015) as does SNP-based heritability estimation (Yang et al. 2015). A possible resolution of this discrepancy is that SNP effect sizes should be drawn from a Poisson distribution rather than a uniform distribution, increasing the incidence of large effect SNPs. Alternatively, in the presence of closely linked antagonistic effects, the 2:2 scenario results suggest that secondary associations will appear to make a smaller contribution to the variance at a locus due to interference.

Another source of mis-estimation of effects is the existence of relatedness among samples. We corrected for this by fitting a GRM in the mixture modeling framework of GEMMA (Zhou and Stephens 2012), a method that simulation confirms is quite effective, although again biased. Independently, we estimated the effect of family structure in the CAGE dataset by comparing the eQTL profiles in CAGE with and without the 800 individuals from the BSGS twin study from Brisbane. The reduced dataset was also analyzed by sequential conditional analysis, but in a straight-forward linear fixed effects model. Due to the larger sample size of the full dataset, an additional 850 eGenes were discovered, while 88% of the loci eGenes discovered in the smaller dataset were retained (*r*^2^>=0.3). The few that were lost were close to the significance threshold initially, so may have experienced sampling boost analogous to the winner’s curse.

Finally, the comparison of the CAGE and FHS datasets allowed us to address the repeatability of multi-SNP eQTL discovery. While the overall estimates of the proportion and identities of eGenes and the incidence of multiple-eSNP loci, is similar in the two datasets, the actual overlap in shared primary association signals is only ~41% percent, and overall just ~29%. The method used to normalize gene expression data can alter eGene discovery for as many as a quarter of all loci (Zeller et al. 2010), and sample size differences also will have influences the replication rate. For the CAGE cohort, the first 10 PCs were fit to control for batch and unknown confounding factors, while for the FHS, 62 significant surrogate variables were removed. To evaluate the influence of these transformations, we also evaluated the eQTL profiles for a subset of the FHS data and found that in moving from fitting 1 to 62 surrogate variables (Leek et al. 2012), there was a consistent trend to discover more eGenes but for these to have a lower replication rate and hence less confidence (Table S4). We also assessed the concordance of estimates for multiple probes per locus, reasoning that some of the differences may reflect regulation either of alternative splicing or exon-specific RNA degradation rates. In the FHS offspring samples (N=2,119), there was a tendency for the probesets that mostly overlap the same exon as the Illumina probe in CAGE, to have a higher replication rate (47.2% with *r*^2^>=0.8). However, within loci there was considerable variation in primary and secondary signals across probes, which we interpret as implying that deconvolution of technical and biological sources of error in multi-SNP identification would not be trivial. Instead, we simply note that technological differences between the two hybridization platforms are another large source of error in eQTL estimation, and no doubt RNASeq will have different properties again.

Thus, while large datasets have very good power for detection of complex regulatory contributions for individual genes, there are a host of technical and statistical reasons why fine mapping of causal variants remains a challenge. There are two immediate strong implications of these results. One is that even though the majority of identified eSNPs are expected to map to credible intervals that include the causal variant (Gusev et al. 2014; Finucane et al. 2015), there will also be many instances where incongruence between the statistical interval and chromatin or other functional evidence (Huang et al. 2015) is not to be expected. The causal variant may simply be poorly mapped due to interference among linked functional sites. This effect may also influence the fine mapping of pleiotropic associations (Fortune et al. 2015). The second implication of the high frequency of multi-site regulation is to emphasize caution in using univariate statistical support for an eSNP effect as sufficient evidence that an association between a SNP and a trait is evidence for causation. At a minimum it is imperative that the full spectrum of eSNP effects across the locus be evaluated to confirm that the site is not simply in LD with higher likelihood eSNPs that are not themselves associated with the trait. Experimental validation of individual sites seems warranted in situations where establishment of the identity of the causal variant(s) is desired.

## Data and Methods

### Datasets

We analyze two different blood eQTL data sets. The Consortium for the Architecture of Gene Expression, CAGE dataset consists of Illumina HT12 microarray-based gene expression profiles, as well as whole genome genotype information, from five research studies: the Brisbane Systems Genetics Study (BSGS, N=926) (Powell et al. 2012), Atlanta-based Centre for Health Discovery and Well-Being (CHDWB, N=439) (Wingo and Gibson 2015) and Emory Cardiology Genebank (N=147, Kim et al. 2014), Estonian Genome Centre - University of Tartu (EGCUT) study (N=1065, Schramm et al. 2014), and the Morocco Lifestyle study (N=188, Idaghdour et al. 2010), for a total of 2,765 individuals. IRB approval was obtained for the combination of data into a mega-analysis both by the University of Queensland and for each participating site. The second dataset from the Framingham Heart Study (FHS) (Huan et al. contains two-generation data based on Affymetrix Genechips. A total of 5,075 participants with both genotype and gene expression information from the offspring (N = 2,119) and third-generation (N = 2,956) cohorts were included in this study. Whole blood samples were collected at the eighth examination of the offspring cohort and the second examination of the third-generation cohort. Raw genotype and gene expression were downloaded from dbGAP (phs000007.v25.p9) with IRB approval.

### Simulation studies

Four different simulation studies were conducted. In all cases, we use the terminology uni-site to refer to models where a single causal variant is modeled as a fixed effect, and multi-site where two or more variants are modeled, also as fixed effects. The term multivariate modeling is used for situations where there are two or more dependent variables, whereas in these models we are assessing the joint effects of two or more causal variables. Some models also incorporate mixed effects of covariates such as a genetic relationship matrix.

The first set of simulations evaluated the effect of family structure on eQTL estimation. Genotype information for chr 1 of the EUR population in the 1000 Genome project (1000 Genomes Project Consortium 2015) was used as the reference from which three causal variants (snp1, snp2, and snp3) with maf >= 0.01 were chosen as eSNPs, either with low (r^2^=0) or intermediate (*r*^2^=0.3) LD. One hundred iterations with simulated heritability of 0.25 were performed for each scenario, simulating family structure with SeqSIMLA2 (Chung et al. 2015), and a sample size of 1,500 (after randomly choosing half of the individuals from each of 250 families, each with two grandparents, four parents, and two-by-three children). Mixed linear models were run in GEMMA (Zhou and Stephens 2012) incorporating a genotypic relationship matrix to account for familial relatedness.

The second simulation study asked whether unimputed variants influence the localization of eSNP signals. Since non-imputed SNPs are not present in the CAGE data, we approximated their identities by choosing randomly sampling from a set of CAGE imputed SNPs weighted to have the same frequency distribution shown in Fig 5C and assigning effect sizes from 2% to 10% of the variance explained for normally distributed pseudo-gene expression traits using the CAGE (minus BSGS) genotypes. We then removed the SNP and all other SNPs with r^2^>0.8, and performed stepwise conditional regression, documenting instances of primary and secondary signals at p<10-5, as plotted in Fig 5E. The cumulative proportion of spurious secondary signals was computed by summing the detection rate by the size of the minor allele frequency bin of the unimputed SNPs.

The third set of simulations assessed the power and accuracy of two-site regressions assuming that the identities of the two causal variants are already known. We modeled the influences of effect size, minor allele frequency (maf), Linkage Disequilibrium (LD), and sample size. Environmental variance was randomly generated as a z-score (mean 0 and standard deviation 1) and genotype effects (β) were added in standard deviation units multiplied by plus or minus 0, 1, or 2 according to genotype so as to account for from 2% to 30% of the phenotypic variance, computed as 2*p*(1-*p*)β^2^. Thus, an allele with β = 0.8 is expected to explain 20% of the variance if maf p = 0.2, or 32% of the variance if p = 0.5. The influence of LD was assessed at *r*^2^ = 0.1, 0.5, or 0.9, noting that as LD increases, high *r*^2^ values are not obtained for combinations of a rare and a common allele. For each combination of parameter values, we generated 1000 randomizations of the environmental variance, and assessed (i) the univariate estimate at each genotype, (ii) the mean conditional estimate of the second SNP, and (iii) the joint effect estimates with both SNPs. From these values we computed the mean absolute value of the deviation between the observed estimate and the true effect size from the univariate, conditional, and joint (two-site) models. The univariate estimates agree extremely well with expectations from the analytical solution described in the results section 3.1.

The fourth set of simulations were performed to evaluate the difference in effect size estimates using the multisite linear regression method for parameter estimation from data representative of the LD structure in the CAGE dataset. For each of 500,000 iterations, 4 sites were chosen at random from a window extending from 200 kb upstream of the transcription start site to 200 kb downstream of the transcription termination site of a randomly picked gene in the CAGE cohort (excluding the BSGS data, since it includes twins), and assigned an effect size from a uniform distribution of variance explained (VE) relative to environmental noise ranging between 0.02 and 0.1. The effect size β for an allele with mafp is computed as V[VE/2p(l-p)]. Subsequently, each phenotype was simulated as 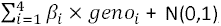, where β_*i*_ is the simulated allelic effect size for a SNP *i*, and *geno*_*i*_ is the dosage of minor allele at the simulated SNP for a given sample. The significance threshold for sequential conditional detection of the variants in a sample of 1,839 CAGE individuals was set at P < 10^−5^, since simulations indicated a <1% FDR for null variants at this level. We evaluated (i) how many of the 4 SNPs were significant in sequential conditional modeling, (ii) the mean LD between each SNP and the other 3 SNPs in the model, (iii) the effect size estimates from the conditional and joint multi-site models, (iv) the difference between these two estimates as a function of the mean LD, and (v) the rank of the discovered SNP for each peak eSNP and the modeled sites, which were assumed to be the causal variants for some trait.

### Genotype imputation

Genotype imputation for the CAGE cohort was performed jointly on the five contributing studies to ensure uniformity of assignment of strand identities of SNPs, and is described in detail in Lloyd-Jones et al. (2016) and at https://github.com/CNSGenomics/impute-pipe. Briefly, the pipeline was as follows: 1. Pre-imputation quality control, and data-consistency checks; 2. Imputation to the 1000G reference panel with Impute2 (Howie et al. 2012); 3. Post-imputation quality control (filtering on various data features); 4. Merging datasets on common SNPs.

For the Framingham Heart Study data, there were a total of 6,950 individuals before imputation. From this set, 29 individuals with genotype missing rate >=5% were removed, and subsequently any SNPs with genotype missing rate >=5% were also removed along with SNPs with Hardy-Weinberg test P <=10^−6^. Prior to imputation, the genotypes were pre-phased using shapeit2 (Delaneau et al. 2013) using the “duohmm” parameter to account for pedigree information. Each chromosome was divided into 5Mb chunks, incorporating the centromere-adjacent region (acen region) into the neighboring chunk, and similarly joining any chunk with <200 SNPs into a neighboring chunk. Imputation was performed with Impute2 (Howie et al. 2012), qctool was used to convert gprobs to gen file format, and only SNPs with info value larger than 0.3 were retained for subsequent analyses. The gen file was converted to plink file format, and SNPs with multiple information and InDel variants were filtered, leaving a total of 8.3 million SNPs with maf greater than 0.01. These were further reduced to ~6 million SNPs with >95% call rate across all 5,075 individuals represented by both genotype and gene expression data.

### Probe Re-annotation

Since SNP imputation for the CAGE cohort was based on hg19/GRCH37, whereas the Illumina probe annotation was based on hg18/GRCH36, we re-annotated the probe information by mapping the probe sequences to hg19/GRCH37 with BWA (Li and Durbin 2009), retaining only the uniquely mapped probes. All probe sequences were secondarily mapped to the reference genome with BLAT (Kent 2002), and only probe sequences uniquely mapped with both methods were determined to be high confidence and subsequently used for eQTL detection. Of a total of 45,931 probes mapped to the reference genome or exome, 7,349 probe sequences mapped to multiple regions or remained unmapped, leaving 38,582 probes taken forward for the eQTL analyses. See Table S1 for summary statistics. Since it is well-known (Walter et al. 2007) that SNPs in a probe influence microarray hybridization, we also discarded 3,856 Illumina probes containing SNPs with maf >1% in the 1000 Genomes European sample (Lappalainen et al. 2013). Similarly, SNPs in the Affymetrix probesets were also converted to positions in the hg19 assembly by applying liftOver (UCSC Genome Browser) to the GPL5188 annotation file downloaded from dbGAP, and annotated to the 1000 Genomes. Any SNP with a maf >1% in the 1000 Genome European population and located within a probeset was deemed to be potentially unreliable, and was included as a covariate during the eQTL estimation steps. Among the 280,000 core probesets, 35,000 have such SNPs, and 15,368 transcripts contain at least one SNP in a probe.

### Gene Expression Normalization

The gene expression normalization strategy for CAGE required aggressive procedures to account for study-specific biases and are described in detail in Lloyd-Jones LR et al. (2016). It consisted of 5 steps: (1) Variance stabilization using the vsn package (Lin et al. 2008); (2) Quantile normalization forcing the intensity distribution across all probes to have the same shape for all samples; (3) Batch effect correction via linear regression to account for known technical effects, such as RNA extraction date, and physical batch; (4) Batch effect correction (via principal component analysis, removing the first 10 PC) to account for unknown confounding procedural, or population-based influences; and (5) Rank normal transformation, namely a final transformation of each probe to a normal distribution with mean 0 and variance 1.

For the FHS data, raw gene expression processed by Affymetrix APT software (version 1.12.0) was downloaded from dbGAP, log_2_ transformed, and surrogate variable analysis (SVA) (Leek et al. 2012) was used to remove confounding factors, fitting a total of 62 surrogate variables by a linear regression model. Note that the FHS gene expression study (Huan et al. 2015) reported results of a different normalization that included fitting blood cell counts, which we chose to avoid since a similar procedure was not applied to the CAGE data.

### Multi-site eQTL Detection

For this study, local SNPs were stringently defined as SNPs located within 200 kb upstream or downstream of the gene (defined as the first TSS and last TES listed in the hg19 annotation) containing the probe. Sequential conditional analyses were performed for each probe, and the genes with significant eSNPs were called eGenes. Since both the CAGE and FHS cohorts contain family-based data (the former for a quarter of the samples from the BSGS twin study (Powell et al. 2012); the latter for all participants), a mixed linear model was used for eQTL detection in GEMMA (Zhou and Stephens 2012), which fits a genetic relatedness matrix (GRM) as a covariate alongside fixed genotype effects. The multiTrans tool (Joo et al. 2016), which accounts for family structure, was used to specify a study-wise false discovery rate of 5% for genes with multiple independent eSNPs, which was empirically observed to be approximately P < 10^−5^. After first scanning for evidence of at least one local eSNP at this threshold, the residuals after fitting the sentinel SNP were used for a sequential conditional scan for an independent secondary eSNP. This process was iterated until no more signals were observed below P = 10^−5^. SNPs in high LD with each previously detected signal (*r*^2^>=0.9) were also filtered out of the sequential analyses. The effect sizes of each discovered SNP were recorded as the sequential conditional estimates. Subsequently, for the multi-site effect size estimates, all discovered independent peak SNPs were fit with the GRM in one mixed model. However, since the GEMMA software does not report the effect sizes of all fixed effects simultaneously, we fit the multi-site models with one SNP specified as the target effect, including the other significant SNPs, as well as the GRM, as covariates. This estimation procedure was repeated for each included SNP, recording the effect size of the target SNP as the multi-site effect, noting that the amount of variance explained by each gene’s models is the same and the effect sizes should be the same. To control the influence of SNPs located in probes in the FHS data, we incorporated in-probe SNPs with an LD *r*^2^ cutoff 0.75 as covariates during the multi-site modeling step. When the genotype of an in-probe SNP was not available, it was imputed from the 1000G data.

## Supplementary Figure legends

<u>Figure S1.</u> Summary of datasets and analytical workflow, including pointers to location of results.

<u>Figure S2.</u> Simulation in the presence of familial relatedness. GEMMA was used to simulate mapping of three eSNPs at a locus under two scenarios: the first assumed no LD, and three causal variants with mafs 0.17, 0.12, and 0.17, assigned beta values 0.46, 0.38, and 0.33 respectively (hence explaining 6%, 3%, and 3% of the variance); the second introduced moderate LD (*r*^2^ = 0.3) between the second and third SNPs, with assumed mafs 0.17, 0.12, and 0.02, and the β values 0.46, 0.38, and 0.87, again for 6%, 3%, and 3% variance explained. Sequential conditional analysis performed on just the three SNPs led to estimation of the proportion of variance explained from the sequential univariate estimate (left plot of each pair), or from the multi-site joint fitting (right plots). Blue dots indicate the true effect size, while box and whiskers show the median, interquartile range, and 95% confidence intervals of estimates from 100 iterations. In both the no LD (A) and intermediate LD (B) scenarios, the multisite model provides more accurate estimates.

<u>Figure S3.</u> Comparison of Framingham and CAGE studies. (A) Percent variance explained by the sum of effects at each of the discovered independent eQTL at each locus measured in the FHS (x-axis) and CAGE (y-axis) datasets. (B) Replication rate of all FHS eSNPs in the CAGE cohort, expressed as the proportion of FHS eSNPs captured by a SNP with LD *r*^2^ > 0.8 in CAGE, in bins of NLP units. (C-F) Replication rates for 5,597 primary, 2,098 secondary, 656 tertiary and 337 quaternary signals.

<u>Figure S4.</u> Simulation of the influence of minor allele frequency and β on allelic effect size estimation. Left side, sequential conditional modeling; right side joint multi-site modeling, with n= 2,000 and (top) LD *r*^2^ = 0.5, (middle) LD *r*^2^ = 0.1 and (bottom) LD *r*^2^ = 0.9. The 25 sectors on each panel show results for combinations of two alleles with maf from 0.1 to 0.5 (left to right, top to bottom), within which each circle represents the average of 100 replicate simulations for allelic effects explaining from 2% to 30% of the expression variance (left to right, bottom to top). Circle size and color is proportional to the absolute value of the deviation between the estimated and actual effect size β in standard deviation units. Transition from white to red represents greater deviation for allele 1; larger circles represent greater deviation from the simulated β for allele 2. Empty fields arise because the indicated level of LD is not possible for the corresponding allele frequencies. Results at varying levels of LD show that sequential univariate biases are particularly strong in the presence of high LD (bottom left), but less pronounced for low LD (midddle left). Missing panels are because high *r*^2^ is not possible for the indicated combinations of maf.

<u>Figure S5.</u> Proportion of variance explained by detected eSNPs in simulations with 2 and 3 causal variants. Box and whiskers show median, inter-quartile range, and 95% confidence limits for the proportion of variance explained under two scenarios. For each panel, from left to right in each simulation, Simu is the variance explained by the known sites, Multi is the result fitting discovered eSNPs jointly, Uni is the result of summing the effects from sequential conditional modeling, and Single is the effect of the peak detected eSNP. Y-axis is the proportion of variance explained.

<u>Figure S6.</u> Effect size estimation bias under the three scenarios with 4:0, 3:1, or 2:2 directions of eSNP effects. Panels from left to right in each row show the average absolute value of the difference between the estimated multisite β and true β, the absolute value of the difference between the estimated sequential conditional β and true β, and the average difference between these values. Each scenario includes 500,000 simulations as in Figure 4.

<u>Figure S7.</u> Effect size estimation bias under the two scenarios with 2 and 3 causal variants. Each scenario includes 90,000 simulations and panels from left to right in each row show the average absolute value of the difference between the estimated multisite β and true β, the absolute value of the difference between the estimated sequential conditional β and true β, and the average difference between these values. (A) Top row is results for two sites where the minor alleles have the same sign of effect, bottom row for two sites with the same sign. (B) Top row is for three sites, all with the same sign, and bottom row is one sites operating in the opposite direction.

<u>Figure S8.</u> Signal ranks of simulated causal variants. The number of simulations in which a discovered variant was the indicated rank (left to right) for the first through fourth discovered variant (front to back). Rank refers to the number of SNPs with a smaller p-value than the modeled causal variant, where 1 implies the causal and discovered are the same SNP, 2 that one other SNP in the LD region had a smaller p-value, and so forth. Only cases where the discovered variant was within *r*^2^ > 0.8 of the causal variant are shown.

**Table S1.** Summary of Illumina probe re-annotation

| Detected | Probes | Mismatch # |
| --- | --- | --- |
| HT12v3 | 48,803 |  |
| Exome | 29,452 | 0 |
|  | 990 | 1 |
|  | 316 | 2 |
| Genome | 13,287 | 0 |
|  | 1,486 | 1 |
|  | 400 | 2 |
| Merge | 45,931 |  |

<sup>#</sup> The number of probes with 0, 1, or 2 mismatches relative to the reference human genome

**<u>Table S2.</u>** Multisite effect estimates in the CAGE and FHS datasets (Online Excel Tables)

**Table S3.**
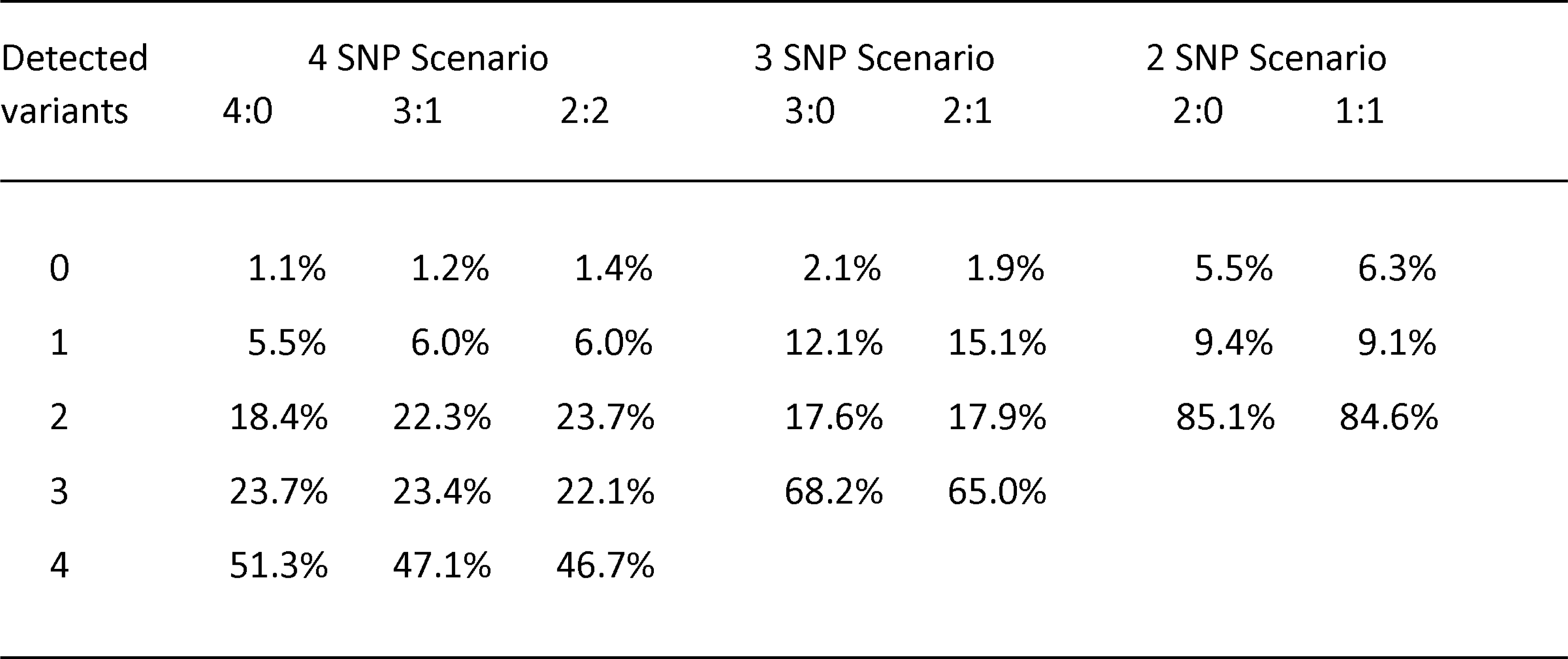
Detected causal variants in simulations with 2 and 3 causal variants

**Table S4.** Effect of fitting principal components on eQTL detection in the FHS

| PCs removed | Explored Genes | eGenes | eQTL | Replicate | Replicate ($r^2 > 0.8$) | Replication Rate |
| --- | --- | --- | --- | --- | --- | --- |
| 1 | 1000 | 389 | 599 | 403 | 154 | 0.38 |
| 3 | 1000 | 429 | 706 | 487 | 166 | 0.34 |
| 10 | 1000 | 462 | 847 | 588 | 185 | 0.31 |
| 14 | 1000 | 468 | 894 | 624 | 195 | 0.31 |
| 18 | 1000 | 479 | 936 | 647 | 195 | 0.30 |
| 24 | 1000 | 489 | 956 | 659 | 193 | 0.29 |
| 30 | 1000 | 492 | 985 | 680 | 192 | 0.28 |
| 32 | 1000 | 496 | 1010 | 700 | 193 | 0.28 |
| 34 | 1000 | 498 | 1009 | 700 | 191 | 0.27 |
| 36 | 1000 | 498 | 1008 | 696 | 189 | 0.27 |
| 44 | 1000 | 502 | 1030 | 708 | 198 | 0.28 |
| 46 | 1000 | 499 | 1035 | 710 | 198 | 0.28 |
| 48 | 1000 | 499 | 1034 | 713 | 199 | 0.28 |

